# Compounds enhancing human sperm motility identified using a high-throughput phenotypic screening platform

**DOI:** 10.1101/2021.09.14.460292

**Authors:** Franz S. Gruber, Zoe C. Johnston, Neil R. Norcross, Irene Georgiou, Caroline Wilson, Kevin D. Read, Ian H. Gilbert, Jason R. Swedlow, Sarah Martins de Silva, Christopher LR Barratt

## Abstract

**Study question:** Can a high-throughput screening platform facilitate male fertility drug discovery?

**Summary answer:** A high-throughput screening platform identified a large number of compounds that enhanced sperm motility.

**What is known already:** Several efforts to find small molecules modulating sperm function have been performed but not using high-throughput technology.

**Study design, size, duration:** Healthy donor semen samples were used and samples were pooled (3-5 donors per pool). Primary screening was performed in singlicate; dose-response screening was performed in duplicate (independent donor pools).

**Participants/materials, setting, methods:** Spermatozoa isolated from healthy donors were prepared by density gradient centrifugation and incubated in 384-well plates with compounds (6.25 uM) to identify those compounds with enhancing effects on motility. A total of ∼17,000 compounds from the following libraries: ReFRAME, Prestwick, Tocris, LOPAC, CLOUD and MMV Pathogen Box were screened. Dose response experiments of screening hits were performed to confirm the enhancing effect on sperm motility. Experiments were performed in a University setting.

**Main results and the role of chance:** From our primary single concentration screening, 105 compounds elicited an enhancing effect on sperm motility compared to DMSO treated wells. Confirmed enhancing compounds were grouped based on their annotated targets/target classes. A major target class, phosphodiesterase inhibitors, were identified in particular PDE10A inhibitors as well as number of compounds not previously identified/known to enhance human sperm motility such as those related to GABA signaling.

**Limitations, reasons for caution:** Compounds have been tested with prepared donor spermatozoa and only incubated for a short period of time. Therefore, the effect of compounds on whole semen or with longer incubation time may be different. All experiments were performed in vitro.

**Wider implications of the findings:** This phenotypic screening assay identified a large number of compounds that increased sperm motility. In addition to furthering our understanding of human sperm function, for example identifying new avenues for discovery, we highlight potential inhibitors as promising start-point for a medicinal chemistry programme for potential enhancement of male infertility. Moreover, with disclosure of the results of screening we present a substantial resource to inform further work in the field

**Study funding/competing interest(s):** This study was supported by the Bill and Melinda Gates Foundation and Scottish Funding Council and Scottish Universities Life Science Alliance.

## Introduction

Sperm dysfunction is the single most common cause of infertility. However, there is an absence of new diagnostic tools and non-MAR (Medically Assisted Reproduction) based treatments for the sub fertile man (Barratt et al., 2017; De Jonge and Barratt, 2019). A fundamental obstacle is the relative paucity of knowledge of the production, formation, maturation and physiological functions of both the normal and dysfunctional spermatozoon. There is an urgent need to address this knowledge gap to formulate appropriate diagnostic assays, develop rational therapies and understand how external factors, such as the environment influence these processes (Barratt et al., 2021).

Although some progress has been made in our understanding of the workings of the mature spermatozoon using tools such as proteomics, electrophysiology and imaging, one area in which there has been minimal progress is the development of an effective high-throughput screening System (HTS) using motile human spermatozoa (Martins da Silva et al., 2017). Current methods for assessment of sperm quality are time-consuming and inappropriate for high-throughput drug discovery (Schiffer et al., 2014; Tardif et al., 2014). One way around this has been to utilize HTS assays with surrogate measures of sperm function such as intracellular calcium concentration [Ca^2+^]_i_ (Martins da Silva et al., 2017; Schiffer et al., 2014). Though informative, these do not directly assess cell function and have key limitations. For example, if [Ca^2+^]_i_ is used as a surrogate of sperm motility, a number of compounds generate an increase in [Ca^2+^]_i_ but have no significant effect on motility (Martins da Silva et al., 2017; McBrinn et al., 2019). To provide a transformative leap in understanding a HTS system for the assessment of quantitative motility is necessary. Recently, a phenotypic platform has been developed which examined human sperm motility in a high throughput manner (Gruber et al., 2020).

Although this HTS system has been used to identify potential compounds for male contraception, i.e. those having a negative effect on human sperm function, it can conversely be used to uncover elements of sperm cell biology and function, and to identify compounds that enhance sperm function. For example, it allows for large scale screening of not only approved drugs, target-class specific libraries (such as ion channels, kinase inhibitors), but also large libraries of chemically-diverse lead-like small molecules that could provide the start-point for medicinal chemistry. In this study, we utilized this HTS system to examine six libraries incorporating ∼17,000 compounds with the dual aim of furthering our understanding of human sperm function and, generating possible start points for a medicinal chemistry programme for potential enhancement of male infertility.

## Materials and Methods

### Experimental design

We used a HTS screening platform to assess the motility of live human spermatozoa. The platform and its development are described in detail in Gruber et al., 2020, and summarised below in brief. The platform was used to screen six compound libraries for their enhancing effects on motility. Whilst we have developed a screening module for detection of the Acrosome Reaction using flow cytometry (Gruber et al., 2020), this was not examined in this study in order to increase throughput and focus on compounds affecting motility. The HTS system and experimental design are illustrated in Figure 1.

**Figure 1.**
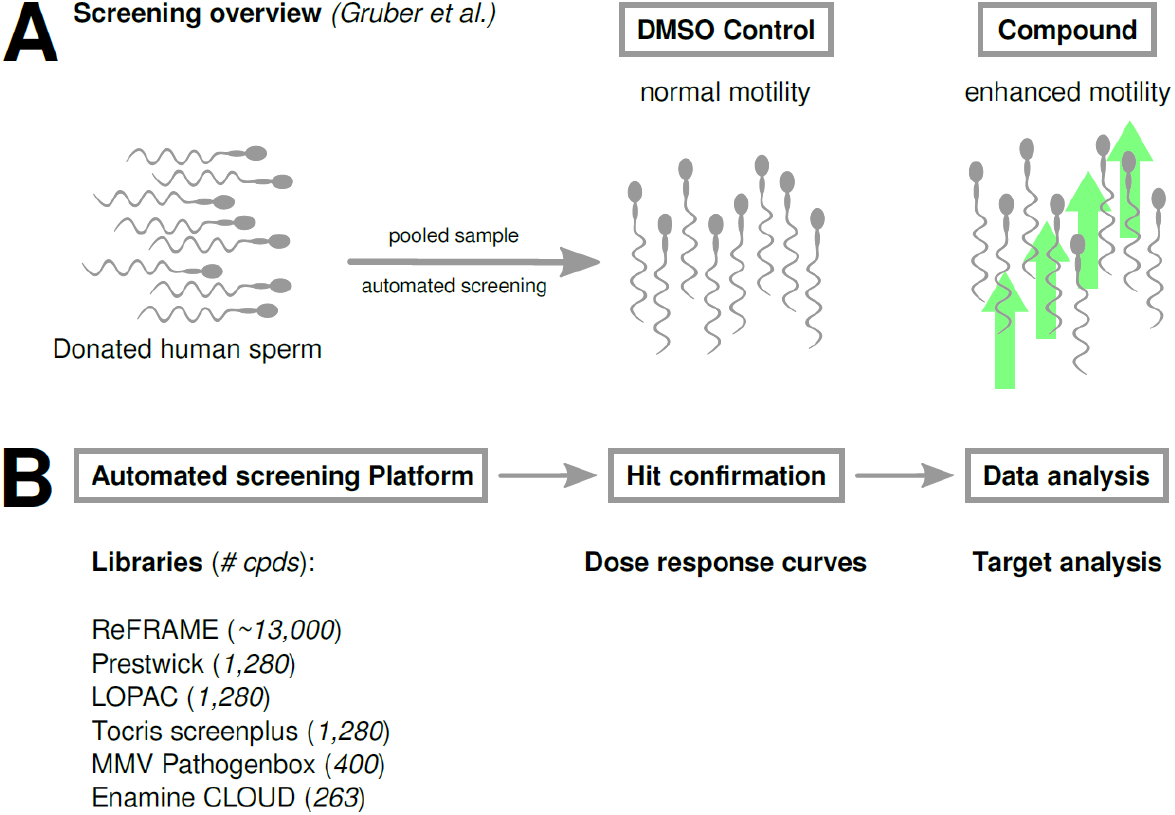
Summary of screening platform and compound screening cascade. **(A)** Motility screening overview as in Gruber et al., 2020. Donated human sperm are pooled and used for automated compound screening to detect compounds which increase sperm motility. DMSO is the vehicle control and the compound label represent a compound which increases motility (reflected by the green arrows). **(B)** Overview of screened compound libraries and follow-up steps. If a compound is selected as a potential hit in the initial screen, dose response experiments are performed (hit confirmation). Analysis of the compounds with confirmed effects by dose response experiment provided some indication of potential target class (Data Analysis).

### Selection and preparation of spermatozoa

Semen samples were obtained from volunteer donors. Written consent was obtained from each donor in accordance with the Human Fertilization and Embryology Authority (HFEA) Code of Practice (version 8) under local ethical approval (13/ES/0091) from the Tayside Committee of Medical Research Ethics B.

Donors had no known fertility problems and normal sperm concentration, motility and semen characteristics according to WHO criteria (2010). Samples were obtained by masturbation, after sexual abstinence of 2-5 days, and delivered to the research laboratory within one hour of production. Samples were allowed to liquify at 37°C for 15 to 30 minutes prior to preparation by discontinuous density gradient centrifugation (DGC). Gradients were prepared using 80% and 40% Percoll (Sigma Aldrich, UK) diluted with non-capacitation media (Minimal Essential Medium Eagle Sigma M3024, supplemented with HEPES, Sodium lactate, and Sodium pyruvate to achieve concentrations previously described (Tardif et al., 2014). For initial screening, prepared donor spermatozoa were routinely pooled to create screening batches of 3-5 donors to reduce donor-to-donor variability. After preparation by DGC and pooling into screening batches, cells were incubated for 3 hours at 37°C under non-capacitating conditions.

### The High-Throughput Screening System

Full details of the HTS system, and its development, are discussed by Gruber and colleagues (Gruber et al., 2020). In brief, screening batches of cells were transferred to a robotic platform (HighRes Biosolutions Inc.) and maintained during the screen at 37°C. Assay-ready 384-well plates, containing compounds were prepared prior to the screen, and filled with approximately 10,000 spermatozoa (20 μL) per well using a liquid handling system (MultiDrop Combi; ThermoFisher). These plates were incubated for 10 minutes prior to imaging. The HTS system utilised a Yokogawa CV7000 Cell Voyager high-throughput microscope to record time-lapse images from 2 positions in each well. An adaptation of a tracking algorithm, Trackpy v0.4.1 (Allan et al., 2018) was utilised to track individual spermatozoa within each well and obtain kinematic parameters. Within the compound-test plates, DMSO was used as the vehicle control.

### Libraries screened

1. The Pathogen box (Medicines for Malaria Venture [MMV] generously provided by MMV, https://www.mmv.org/mmv-open/pathogen-box/about-pathogen-box.). This is a small repurposing library assembled to screen against rare and neglected tropical diseases containing ∼400 diverse, drug-like molecules with demonstrated biological activity against different pathogens.
2. The CeMM Library of Unique Drugs (CLOUD) purchased from Enamine (https://enamine.net/hit-finding/compound-collections/bioreference-compounds/the-comprehensive-drug-collection-cloud) is a set of 263 small molecules representing the target and chemical space of FDA-approved drugs that has been used for drug repurposing.
3. Tocris compound library: (Tocris, Bristol, United Kingdom, https://www.tocris.com/products/tocriscreen-plus_5840.) comprising 1280 biologically active small molecule compounds.
4. LOPAC^®^1280LOPAC (Library of Pharmacologically Active Compounds https://www.sigmaaldrich.com/life-science/cell-biology/bioactive-small-molecules/lopac1280-navigator.html). These compose a biologically annotated collection of inhibitors, receptor ligands, pharma-developed tools, and approved drugs which impacts many signalling pathways and covers all major drug target classes (1280 compounds).
5. Prestwick Chemical Library: (http://www.prestwickchemical.com/libraries-screening-lib-pcl.html) Comprising 1280 off patent drugs with high chemical and pharmacological diversity as well as known bioavailability and safety in humans (1280 compounds).
6. ReFRAME (Repurposing, Focused Rescue, and Accelerated Medchem). The initial library consists of ∼ 12,000 approved drugs, in-development small molecules and bio-active compounds and was constructed as a library for potential drug repurposing (Janes et al., 2018). The advantages of using this library was the potential to identify chemical compounds that are already approved for other indications or having undergone (or currently undergoing) clinical trials or had IND approval – hence potentially accelerating progress towards a safe and effective enhancer of motility. Calibr kindly provided up to 1% of hits for subsequent dose response experiments. Only structures of confirmed hits were unblinded. We have also received a further 950 cpds, which have been added at a later stage to the ReFRAME collection.

### Data normalization and hit confirmation

All steps were performed as previously described (Gruber et al., 2020). In summary, data from every compound well was normalized to those from in-plate DMSO controls (wells containing the same amount of DMSO as compound wells). Two positions were recorded in every well and the average of those positions was used for calculating % of control = (VCL median/DMSO median) x 100, where VCL median is the median of all sperm tracks in each well (immotile, non-progressively motile, progressively motile) and DMSO median is the median of all 16 DMSO control wells on the corresponding plate. Hits from the primary screening were chosen based on % change compared to the control: 20% (LOPAC, CLOUD, Tocris, Prestwick, ReFRAME last batch); 40% (ReFRAME first batch, MMV Pathogenbox).

Hit compounds were examined in subsequent dose response experiments using different donor sample pools (comprising 3 different biological replicates). A compound was only identified as having a positive effect if it was confirmed in these dose response experiments. Dose response curves (DRC) consisting of a series of 8-points with 3-fold dilution (with 10 µM top concentration) were fitted using the DR4PL (4 parameter logistic fit) package in R from which ECx (half-maximum effective concentration of x % effect) was estimated.

### Hit analysis

Chemical space was visualized by generating Morgan fingerprints using RDKit (radius = 2, bits = 2048) and UMAP (McInnes et al., 2018). Physico-chemical properties were calculated using RDKit (http://www.rdkit.org) and KNIME (Berthold M.R. et al., 2008).

## Results

In order to find compounds that positively enhance sperm motility, we used a HTS platform (Figure 1A), to screen ∼17,000 compounds comprising a variety of small molecule libraries (Figure 1B). Primary hits were identified based on percentage effect relative to DMSO control. These limits varied among the libraries (Table 1) and a hit selection criterion of 20% increase in motility (i.e. 120% of control) gave a primary hit rate of between 0.3-1.9% and several chemical start points for further investigations. Primary hit compounds were confirmed in subsequent dose response experiments (two biological replicates, performed on two different days) and, in total 105 compounds were identified as confirmed hits (Supplementary Table 1), with moderate to high motility enhancing activity (>20-40% increase in sperm motility) (Table 1, Figure 2A). In general, motility enhancing compounds increased the fraction of progressively motile cells (Figure 2B-C). The confirmed hits were annotated (broad definitions based on vendor annotations) to affect a variety of protein target classes (Figure 3A, Supplement Table 1).

**Table 1.** Summary of screened libraries

| Library | No. of compounds | No. of increaser hits | Hit cut-off<sup>2)</sup> | % Hit rate |
| --- | --- | --- | --- | --- |
| ReFRAME | ~13,000 | 48 | 20-40 %<sup>1)</sup> | 0.3 |
| Prestwick Chemical Library | 1,280 | 4 | 20 % | 0.3 |
| Tocriscreen Plus | 1,280 | 24 | 20 % | 1.9 |
| LOPAC | 1,280 | 20 | 20 % | 1.6 |
| MMV Pathogenbox | 400 | 8 | 40 % | 2 |
| CLOUD | 263 | 3 | 20 % | 1.1 |
| **Total** | **17,503** | **107** | **-** | **1.2** |

<sup>1)</sup> max. 1 % resupply <sup>2)</sup> increase relative to DMSO control

**Figure 2:**
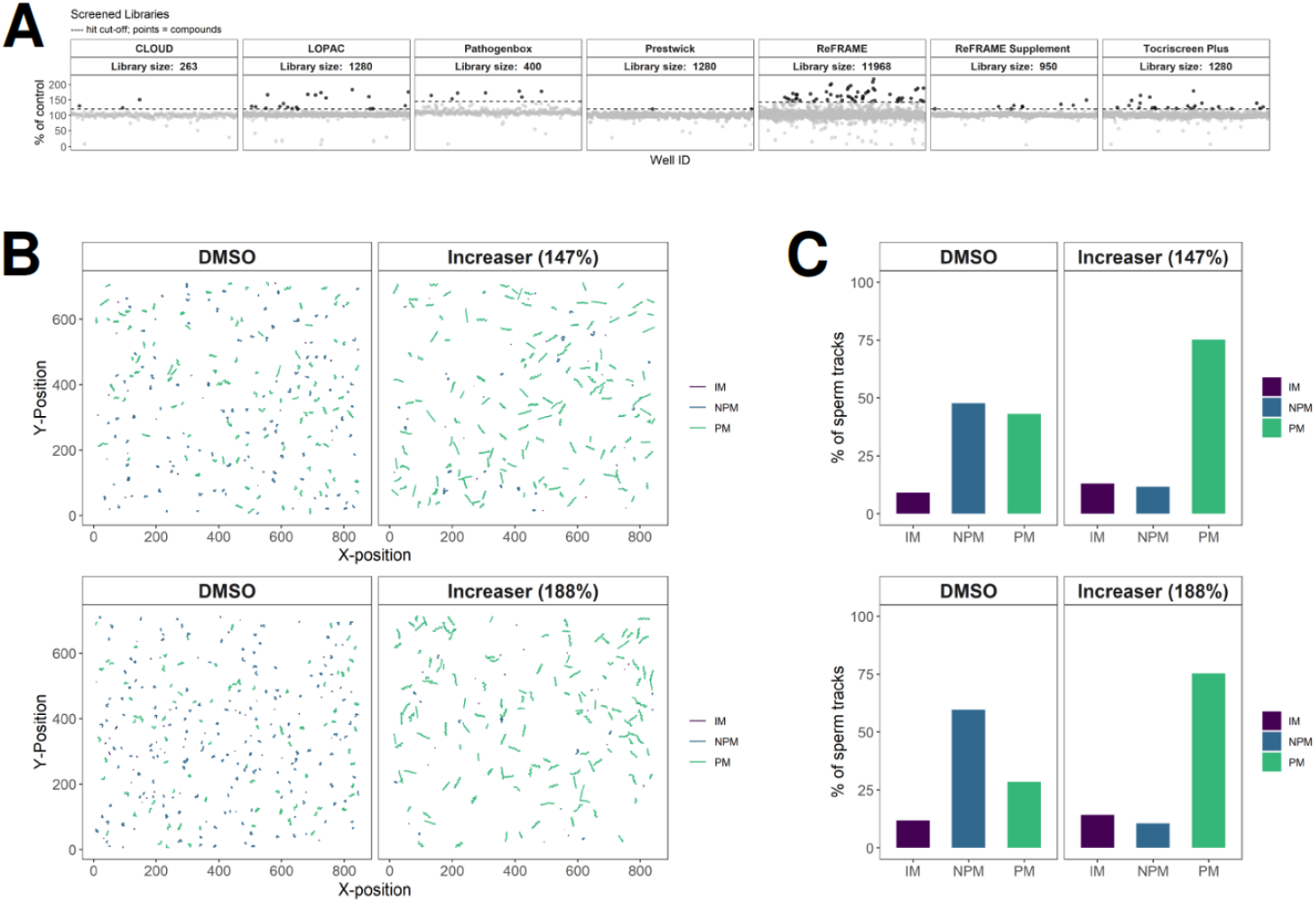
Primary screening results with examples. **(A)** Primary screening data from all screened libraries. Data is presented as % of control, which is defined as a well median of VCL (including all sperm tracks) relative to a median of VCL of vehicle (DMSO) control wells. Dashed lines represent hit cut-offs for each library. Each dot represents and individual compound and black dots represent hits with motility increase above cut-off. **(B)** Example tracks of spermatozoa exposed to two compounds which increased motility to different degrees; ∼147% of control (upper panel) and ∼188% of control (lower panel) against a corresponding DMSO control well. Track colour indicates sperm track classification: purple (IM; immotile), blue (NPM; non-progressively motile), green (PM; progressively motile). **(C)** Quantification of classified sperm tracks of (B).

**Figure 3.**
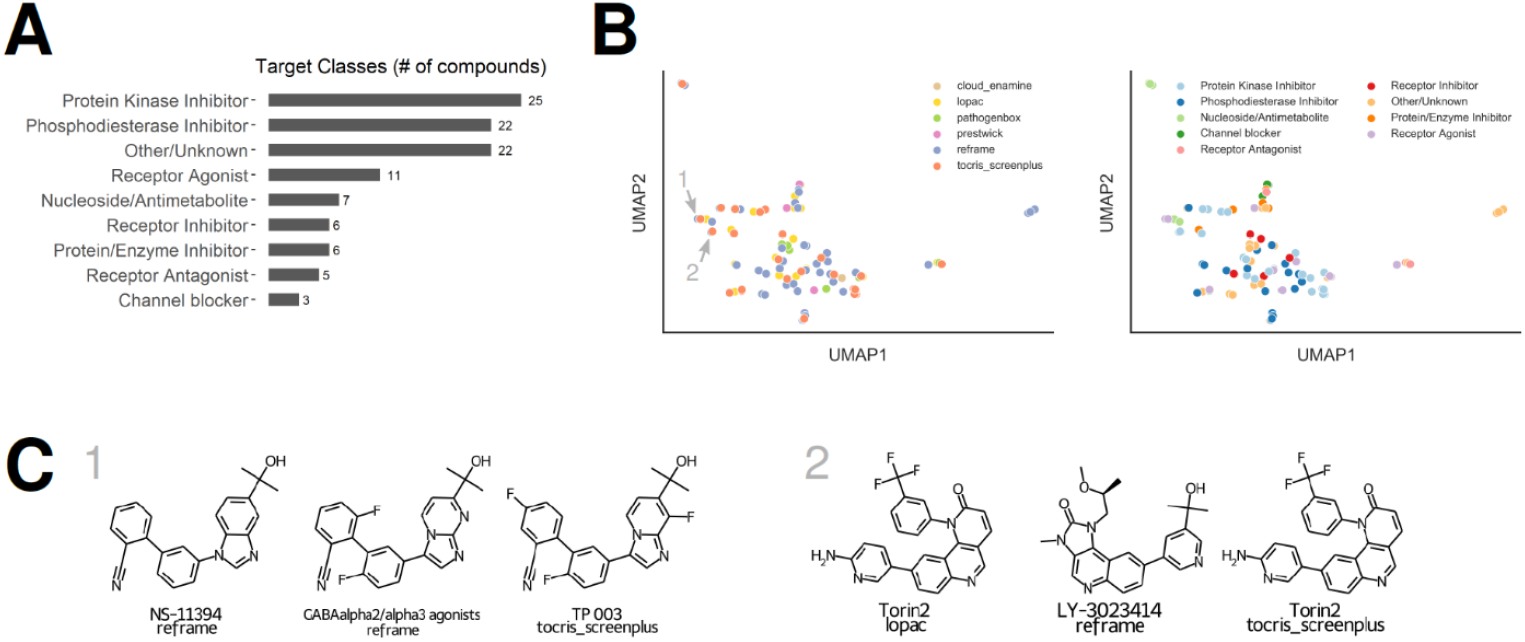
Classification of screening results. **(A)** Summary of the target classes of compounds confirmed by dose response experiments. Target classes were identified according to library annotations. The “other/unknown” category is comprised of compounds with no annotation available from the library vendor or unknown mode of action. **(B)** Chemical space visualization of motility enhancing compounds. Each enhancing compound has been encoded as chemical fingerprint (Morgan Fingerprint) with 2048 bit features. All features have been reduced to two dimensions using UMAP. Color indicates screening library (left panel) or annotated target class (right panel). **(C)** Examples of similar hit structures with names and library information related to GABA signaling (panel 1, also highlighted in B) and mTOR signaling (panel 2, also highlighted in B).

Some compounds could not be assigned to a clear target class, however, of the annotated compounds, protein kinase and phosphodiesterase were the most common target classes (Figure 3A). Another prominent target class were receptor modulators (inhibitor, agonist, antagonist), some of which are related to GABA signalling (Figure 3A, Figure 3C panel 1, Supplementary Table 1).

The most potent hits had sub-micromolar potency and substantial effects on motility (up to 190% compared to DMSO controls Table 2). Chemical space visualization (Figure 3B) reveals that several confirmed hits have similar or identical structures (Figure 3C). The small molecule libraries used had some overlapping compounds and a number of these were either consistently active (e.g. SCH 58261, Torin2 (Figure 3C panel 2), or Ethaverine) or consistently inactive (e.g. Rolipram, Milrinone or Sildenafil). A small number of compounds, for example Papaverine, were active in one library but not others. This may be due to the concentration at which they have an effect and/or inconsistencies with the library material such as purity/quality of compounds.

**Table 2.** Most potent increasing compounds

| Compound | EC<sub>x</sub><br>[μM]<sup>1)</sup> | [% of control] | Target Action |
| --- | --- | --- | --- |
| TAK-063 | 0.04 | 145 | Phosphodiesterase Inhibitor |
| RFM-012-216-7 | 0.14 | 128 | Unknown |
| GW 843682X | 0.17 | 192 | Protein Kinase Inhibitor |
| Torin2 | 0.22 | 196 | Protein Inhibitor |
| Linsitinib | 0.24 | 191 | Receptor Inhibitor |
| Tolafentrine | 0.26 | 153 | Phosphodiesterase Inhibitor |
| Epetirimod | 0.32 | 166 | Unknown |
| E6005 | 0.33 | 151 | Phosphodiesterase Inhibitor |
| GABAα2/α3 agonists | 0.35 | 138 | Receptor Agonist |
| JNJ-42396302 | 0.39 | 158 | Phosphodiesterase Inhibitor |
| NM-702 | 0.48 | 152 | Phosphodiesterase Inhibitor |
| RG-7203 | 0.49 | 162 | Phosphodiesterase Inhibitor |
| Trequinsin hydrochloride | 0.5 | 143 | Phosphodiesterase Inhibitor |
| Dextofisopam | 0.54 | 143 | Unknown |
| Papverine | 0.55 | 175 | Phosphodiesterase Inhibitor |
| LY-3023414 | 0.56 | 140 | Protein Kinase Inhibitor |
| KF 15832 | 0.61 | 143 | Phosphodiesterase Inhibitor |
| STL515575 | 0.62 | 140 | Unknown |
| Carbazeran | 0.63 | 146 | Phosphodiesterase Inhibitor |

<sup>1)</sup> Half maximal effective concentration

One prominent target class of confirmed hits were phosphodiesterase (PDE) inhibitors (Table 2, Figure 3A, Figure 4A). Some of those are well described such as Trequinsin or Papaverine as influencing motility but for others, such as TAK-063, JNJ-42396302, RG-7203 and PF-2545920 (Figure 4B) no data was available. The latter four compounds have an annotation of PDE10A inhibition (Table 3) and showed sub micromolar responses (0.04 – 0.49 µM) in dose response experiments yet some are structurally distinct (Figure 4B). All annotated PDE10A inhibitors used in this study had a enhancing effect on motility. For all four PDE10A inhibitors a similar shift of VCL relative to DMSO controls was observed (Figure 4C). Another interesting observation among PDE inhibitors is that, at 6µM, we did not detect any enhancing effect of compounds belonging to the methylxanthine class e.g. IBMX or Pentoxifylline. All annotated PDE inhibitors used in this study are summarized in Supplementary table 2.

**Figure 4.**
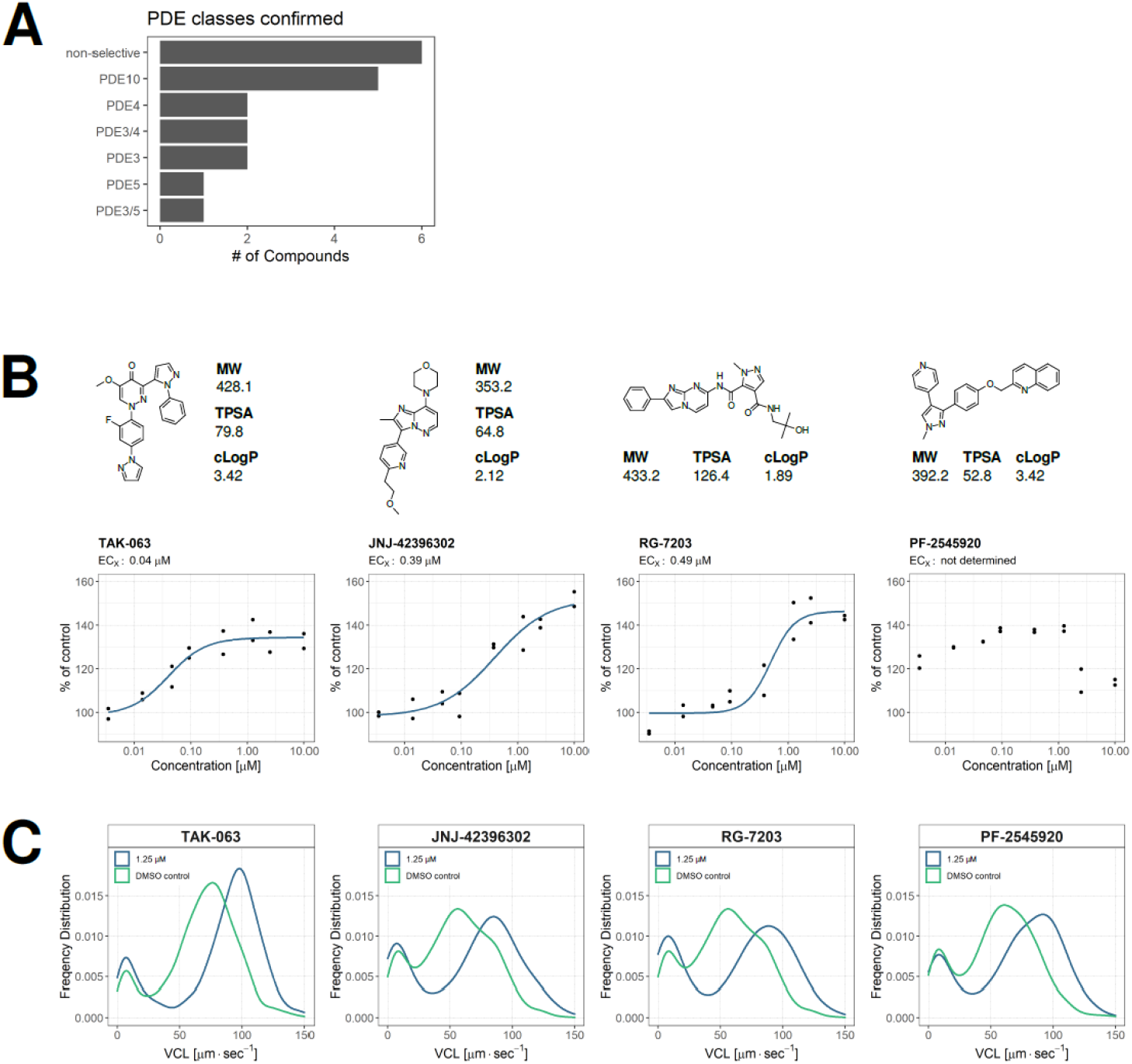
Confirmation of PDE10A inhibitor hits. **(A)** Summary graph of PDE inhibitor classes based on vendor annotation or available information resources (ChEMBL, PubChem, DrugBank). **(B)** Dose response curves of 4 PDA10A inhibitors, with structures and physico-chemical properties. Blue line: 4 parameter logistic model. ECx: estimated half-maximum concentration. Each dot represents an individual data point, n = 2 for each concentration with data collected from two independent dose response experiments utilizing different biological material (i.e. pooled spermatozoa samples from different donors in each experiment). Physico-chemical properties are defined as: MW: molecular weight; TPSA: topological polar surface area; cLogP: computed Crippen-LogP. **(C)** Frequency distributions of sperm VCL of each PDE10A inhibitor shown in (B) at 1.25 µM concentration (blue) compared to DMSO control wells (green).

**Table 3.** PDE10A inhibitors

| Compound | EC<sub>x</sub><br>[μM]<sup>1)</sup> | [% of control] | PDE<br>class |
| --- | --- | --- | --- |
| TAK-063 | 0.04 | 145 | PDE10A |
| JNJ-42396302 | 0.39 | 158 | PDE10A |
| RG-7203 | 0.49 | 162 | PDE10A |
| PF-2545920 | n.d.<sup>2)</sup> | 130 | PDE10A |

<sup>1)</sup> Half maximal effective concentration <sup>2)</sup>EC<sub>x</sub> not determined;

## Discussion

The current study utilised a validated imaging-based screening platform that measures fundamental aspects of human sperm behaviour (Gruber et al., 2020) to screen a collection of chemical libraries comprising ∼17,000 approved drugs, clinically-tested compounds and annotated chemical tool compounds for their potential to enhance motility. The aim was to further our understanding of human sperm function and generate possible start points for a medicinal chemistry programme for potential enhancement of male infertility.

There are significant challenges in producing a suitable platform for HTS of mature human spermatozoa (see(Gruber et al., 2020) and development is always a balance between achieving the necessary high throughput and a detailed assessment of each compound. In these experiments initial screening was performed under non capacitating conditions at one concentration (6µM) with compounds being assessed after only a relatively short incubation (10-27 mins). The data will therefore primarily reflect the use of these conditions and its possible that other permutations, for example, screening under capacitating conditions or longer incubation times may generate different results. In the current study when a primary hit was identified, dose response experiments using different pools of donor cells were undertaken to confirm the hit and provide initial information on potential activity. Although this approach provides information about the activity of the compound, further experiments are necessary to provide a more comprehensive picture of each compound’s activity and assess their mechanisms of action and suitability for further development (see (McBrinn et al., 2019) for examples of such investigations).

The screening platform is complementary to a reductionist approach. Identification of several PDE inhibitors as confirmed hits in dose response experiments (discussed below) provide evidence of the robustness of the HTS platform. Several phosphodiesterase inhibitors (PDEi) (e.g. ibudilast, trequinsin hydrochloride, and papaverine) have previously been shown to significantly increase human sperm motility, confirming the ability of the HTS platform to identify compounds which are effective at or below concentrations of 6μM. Furthermore, identical, or related compounds which were present in two or more libraries were identified. For example, ibudilast was present and detected as a hit in both the LOPAC and TOCRIS libraries, and Trequinsin hydrochloride was confirmed in the ReFRAME and TOCRIS libraries. Another example is Torin2, a small molecule mTOR inhibitor, which was also detected in two libraries (LOPAC and TOCRIS) along with a structurally related compound (LY-3023414, see Figure 3C panel 2) another compound with annotated activity against mTOR. This is intriguing as it has been recently described that in older men mTORC1 is inhibited in highly motile spermatozoa compared to their defective/immotile counterparts (Silva et al., 2019). For the largest library screened, the ReFRAME set, only 1% of the hits were un-blinded, limiting our ability to analyse less active and inactive compounds from this set.

Of the target classes identified, PDE inhibitors (PDEi) account for 18/105 of the compounds found to increase sperm motility. This is not surprising and several of the PDEi hits have been previously identified to increase human sperm motility e.g. Dipyrimadole, Ibudilast, and Papaverine(Tardif et al., 2014). Another potent confirmed hit, trequinsin hydrochloride, has been extensively examined by McBrinn who described the compound’s effects on human sperm motility and function (McBrinn et al., 2019). Strikingly, a proportion of the PDEi hit compounds are annotated as specific to PDE10A. Although relatively little has been published on the effects of PDE10A inhibitors on human sperm, the presence of the active PDE10A enzyme has been confirmed (Marechal et al., 2017). Papaverine, one of our PDE10A inhibitor hit compounds, was one of the specific PDE10A inhibitors have previously been used at high concentrations to mimic the effects of capacitation and increase the progesterone induced calcium response (Torres-Flores et al., 2008). Marechal and colleagues also confirm their findings in additional experiments with the newly available PDE10A inhibitor MP-10 (Marechal et al., 2017). MP-10, also known as Mardepodect or PF-2545920, was a hit in our screen (Supplementary Table 2). Little information is available for the other PDE10A inhibitors. TAK-063 has gained interest as a potential therapeutic drug in the treatment of schizophrenia (Suzuki and Kimura, 2018) and is the subject of clinical trials. While JNJ-42396302 has undergone phase 1 clinical trials, it has, to our knowledge not been previously tested on spermatozoa. The high representation of PDE10A inhibitors in this screen, combined with their apparent potency, could indicate their potential for further investigations for use in infertility treatment and or MAR.

Several PDE inhibitors which have been well documented for their effects on motility parameters of human sperm, including pentoxifylline did not appear as a hit in this screen. Pentoxifylline belongs to the methylxanthine class of drugs which includes aminophylline, theophylline, pentoxifylline, caffeine, and 3-Isobutyl-1-methylxanthine (IBMX). Although all of these drugs were screened, none increased sperm motility above the selection threshold. While this might initially be surprising, it is worth noting that initial screening conditions were at 6 μM for 10-27 minute incubation and the actions of these drugs may require higher doses and/or longer incubation time. IBMX, for example, is used at concentrations from 30 μM to 1 mM (Lefievre et al., 2000; Marechal et al., 2017; Pons-Rejraji et al., 2011; Tardif et al., 2014). Similarly, pentoxifylline has been used at 3 to 4 mM (Burger et al., 2000; Patrizio et al., 2000; Terriou et al., 2000; Tesarik et al., 1992) although conflicting reports have found no improvement in human sperm motility at the same concentrations (Mathieu et al., 1994; Tournaye et al., 1994) and higher concentrations of 10mM have been used to examine its effects on spermatozoa DNA damage (Banihani et al., 2018). Other such PDE inhibiting compounds included Milrinone, a PDE3 inhibitor shown to effect human spermatozoa motility at 50 μM (Lefiévre et al., 2002), and rolipram, a PDE4 inhibitor with effects at 10 µM (Marechal et al., 2017). Sildenafil and its analog Vardenafil were also screened without appearing as a hit. The cGMP specific PDE5 is expressed at low levels in human spermatozoa (Lefiévre et al., 2002) and its inhibition, in vitro, using sildenafil can improve sperm motility. However, conflicting literature has reported that this effect requires vastly different concentrations of the drug. Lefièvre *et al* report that an increase in progressive motilty required concentrations of at least 100μM, while Glenn et al report an improvement in progressive motility with just 0.67 μM (Glenn et al., 2007; Lefievre et al., 2000).

A substantial advantage of phenotypic screening is that it potentially opens new avenues for investigation allowing discovery of new avenues to improve our understanding of cell. In this screen, in addition to those addressed above, there are several examples that warrant further investigation. For instance, enhancement of sperm motility by Linsitinib which selectively inhibits IGF-1R and the insulin receptor, is in keeping with the recent data of insulin modulating human sperm survival (Aitken et al., 2021). Another novel consistent finding was that modulation of γ-Aminobutyric acid – GABA- resulted in an increase in sperm motility (Supplementary table 1). While there is significant literature on the role of GABA in induction of the acrosome reaction there is surprisingly little relating to human sperm motility. In the current data GABAalpha2/alpha3 agonist and NS11394 (a GABA_A_ receptor modulator) significantly increased sperm motility. Both are selective positive allosteric modulators of GABA_A_Rs albeit working on different GABA_A_ receptor subtypes. Usually they are inert in the absence of GABA or equivalent agonist. Moreover, TP003 and U90042, also GABA_A_ receptor agonists were identified. As for the examination of the insulin receptor pathways more detailed experiments are required but modulation of GABA and associated receptor complexes will uncover as yet undetermined biology related to human sperm motility.

In summary, using a novel HTS, we identified a large number of compounds that increased sperm motility. In addition to furthering our understanding of human sperm function, for example identifying new avenues for discovery such as the role of GABA in sperm motility, we highlighted PDE10A inhibitors as promising start-point for a medicinal chemistry programme for potential enhancement of male infertility. Moreover, with full disclosure of the results of screening we present a detailed resource to inform further work in the field (Supplementary table 1)

## Authors’ roles

FSG performed the sperm preparation, HTS screening, processing of the data. FSG, ZCJ and CLRB analysed and interpreted the data. SMDS, IHG and KDR designed the study, assisted with interpretation of the data and original obtained funding. All authors contributed to the construction, writing, and analysis of data. All authors approved the final manuscript.

## Acknowledgements

We are very grateful to all members of the research team for their invaluable assistance. We also want to thank all the volunteer sperm donors who took part in this study and members of the research group for recruitment. We want to thank Dr David Mortimer and Dr Sharon Mortimer for their helpful insights into comparisons with the CASA system. Thanks go to Dr Steve Publicover for critical reading of the manuscript. Thanks are also due to NPSC lab members for help, particularly John Raynor for engineering support. We thank Mitch Hull, Emily Chen and Kelli Kunen at CALIBR for their help in library plating, logistics and supply of ReFRAME data. We would also like to thank Medicines for Malaria Venture (MMV) for providing pathogenbox reagents.

## Availability of data

The data underlying this article are available in the article and in its online supplementary material.

---

**Supplementary Table 1.**
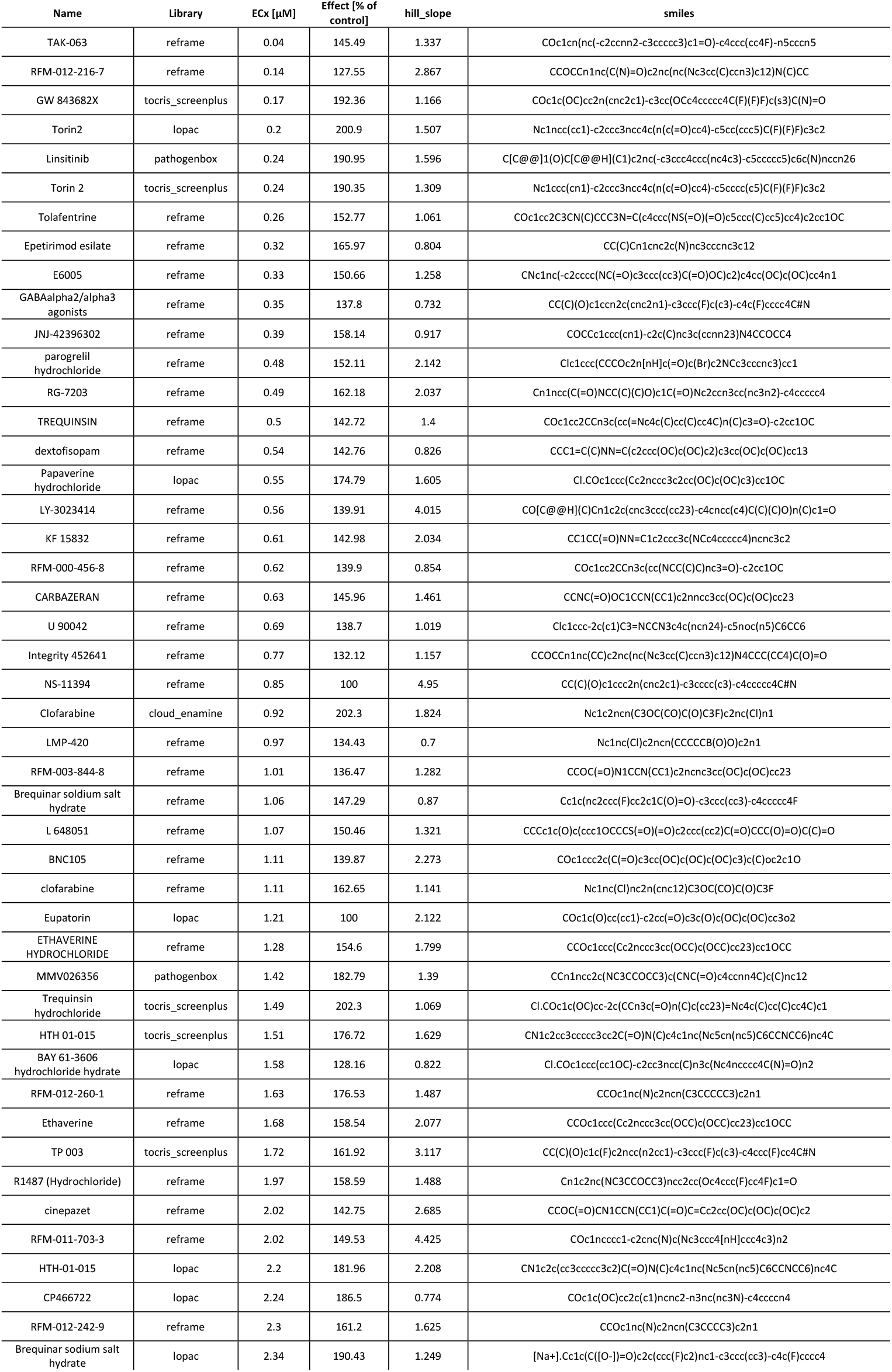
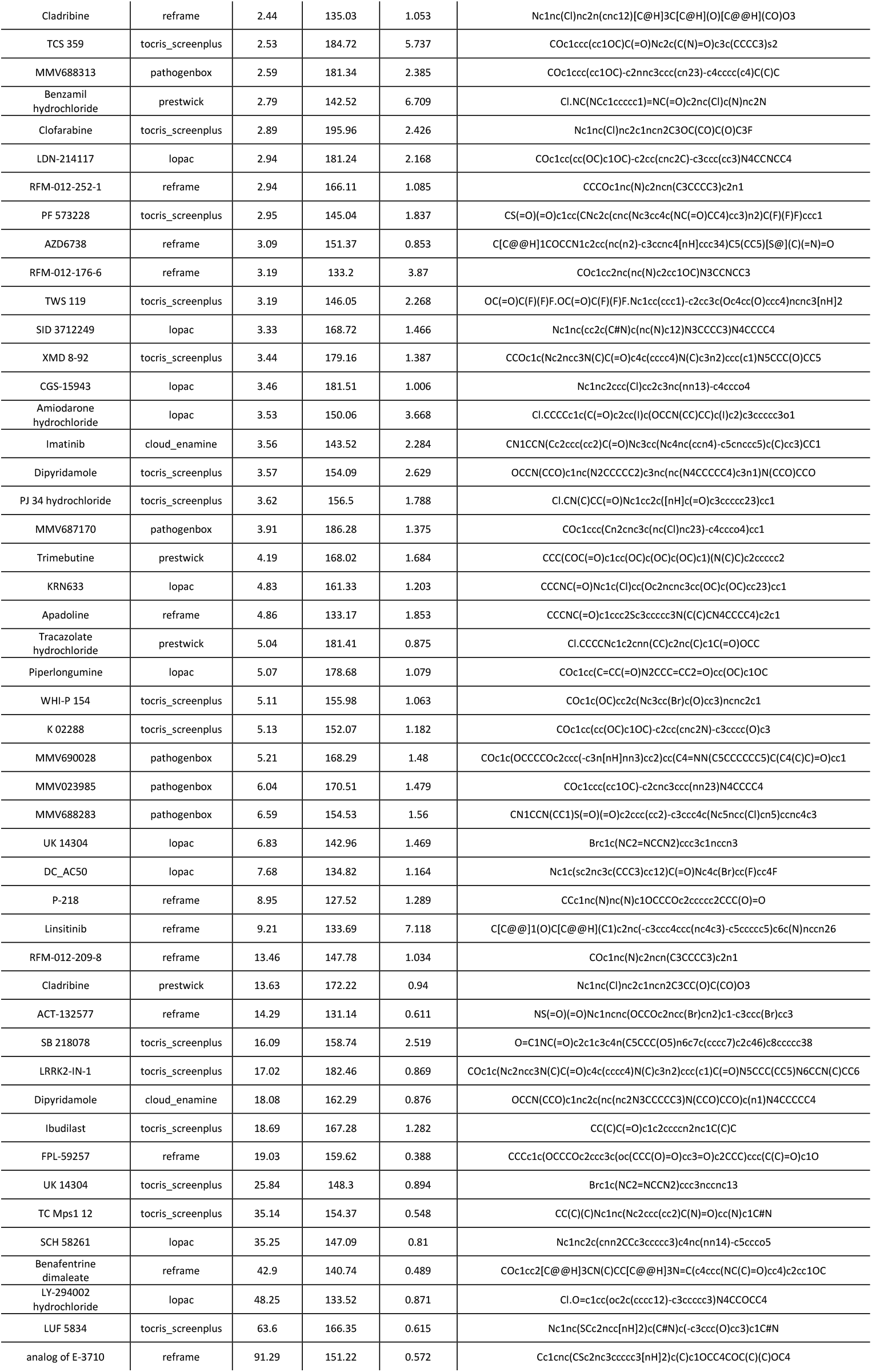
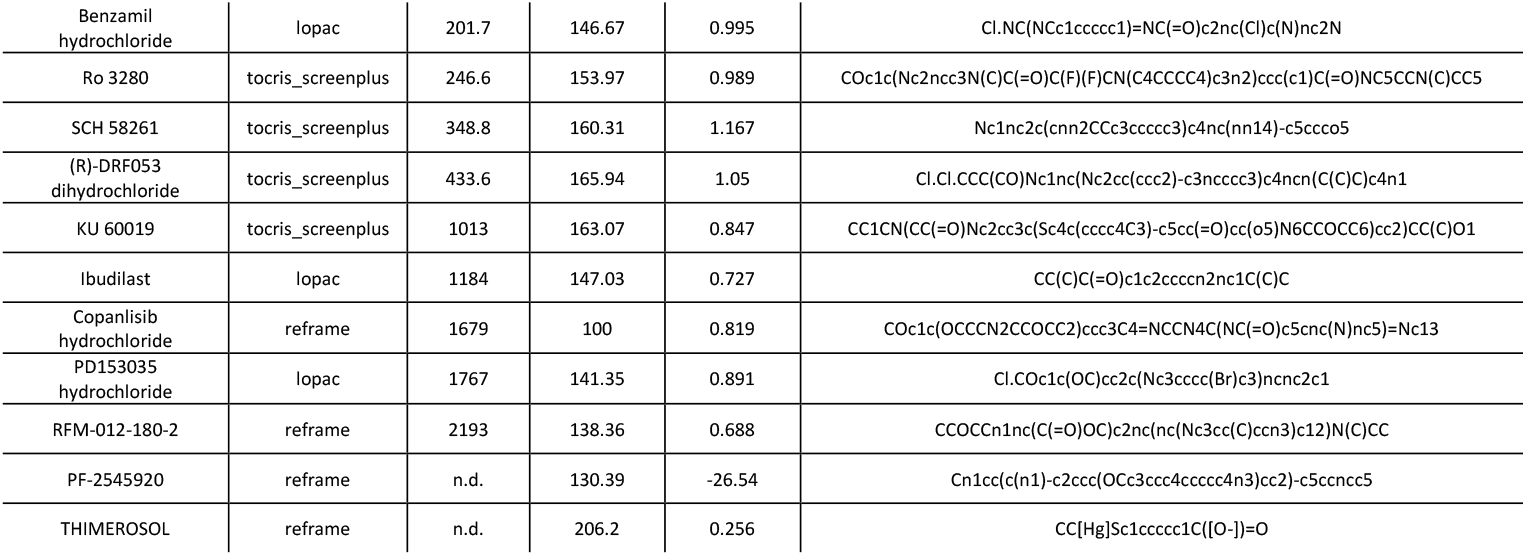

**Supplementary Table 2.**
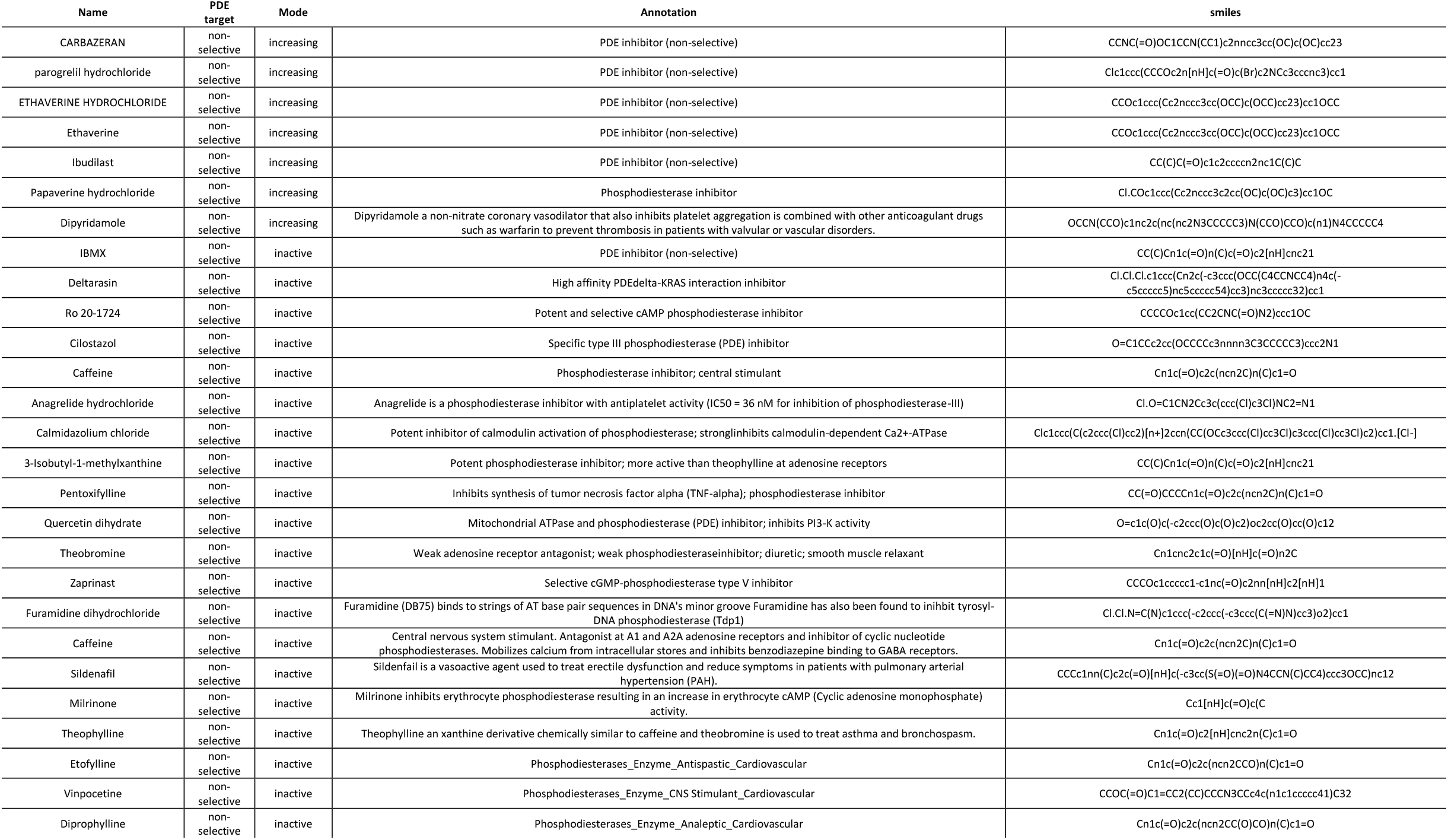
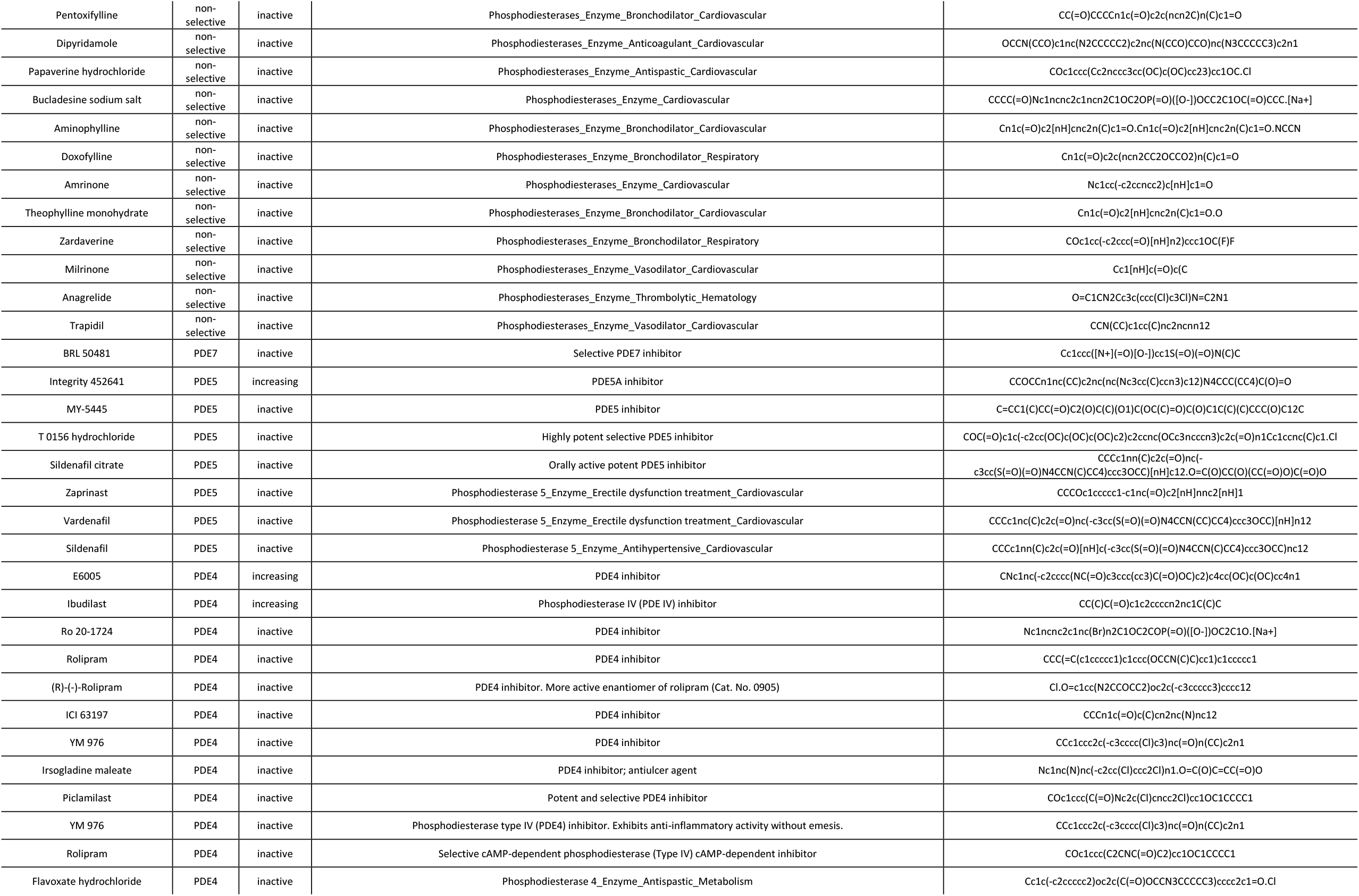
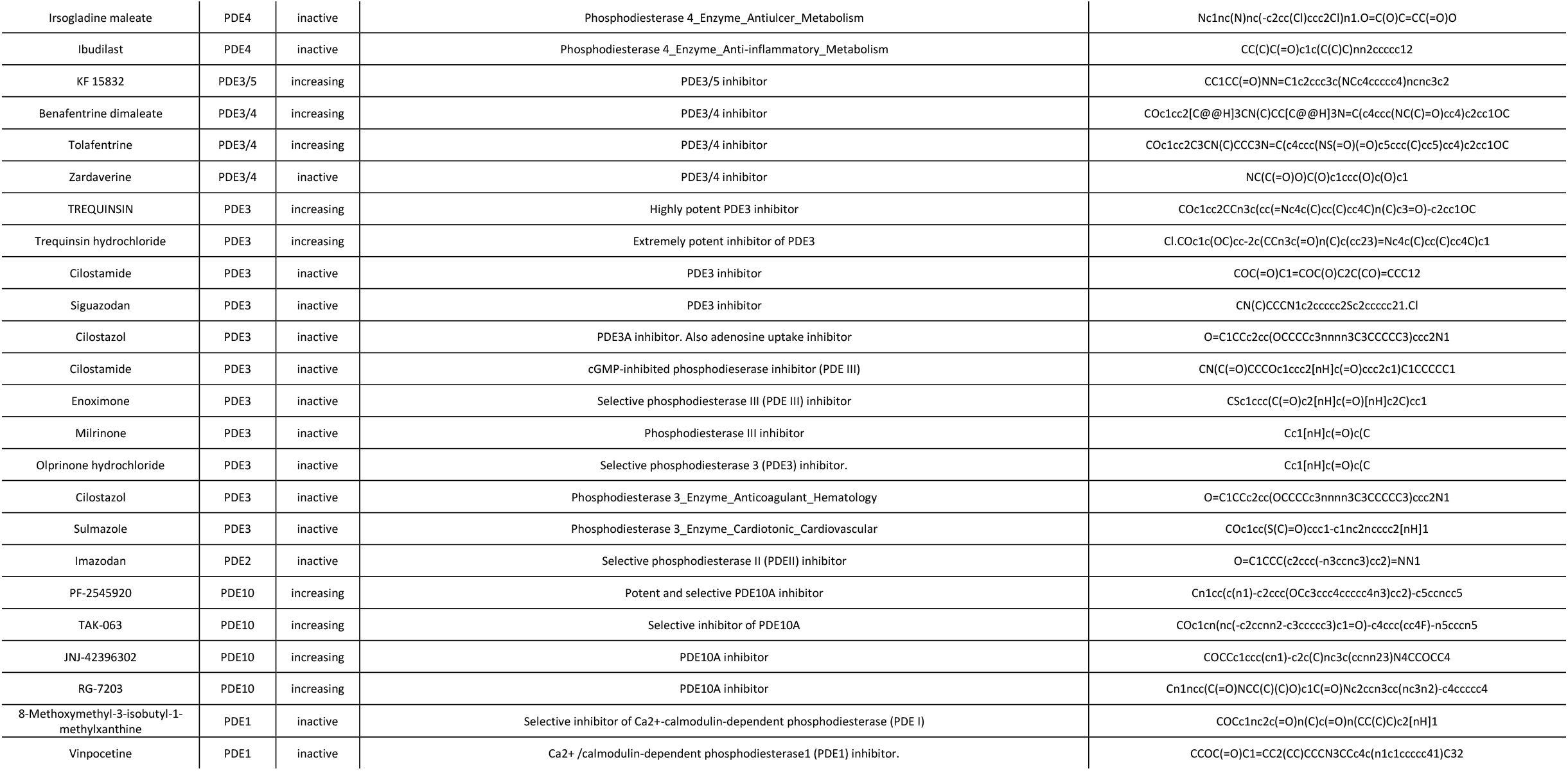

